# Coronin-1 is necessary for enteric pathogen-induced transcytosis across human ileal enteroid monolayers expressing M cells

**DOI:** 10.1101/2020.10.12.305565

**Authors:** Janet F. Staab, Michele Doucet, Rachel Latanich, Sun Lee, Mary K. Estes, James B. Kaper, Nicholas C. Zachos

## Abstract

In the intestine, luminal sampling by microfold (M) cells is crucial for inducing protective mucosal immune responses but can also serve as an entry pathway for pathogens, including bacteria and viruses. Enteric pathogens can influence intestinal M cell function; however, the molecular mechanisms involved in the regulation of uptake and transcytosis of gut cargo by human M cells remain to be determined. Understanding the mechanisms responsible for regulating human M cell function requires a relevant human model. In this study, human ileal enteroids established from healthy donors were grown as confluent monolayers on permeable supports and differentiated to express mature M cells. Enteric pathogens including enteropathogenic *E. coli* (EPEC), adherent invasive E. coli (AIEC), and human rotavirus were apically exposed to M cell enteroid monolayers. M cell-mediated uptake and transcytosis was compared in enteroids infected by pathogenic or commensal bacteria (HS strain). EPEC and AIEC, but not HS, stimulated M cell uptake and transcytosis. We discovered that this pathogen-specific effect was dependent on expression of coronin 1a, a cytoskeletal remodeling protein. Using stable coronin 1a knockdown (KD) enteroids, we observed that EPEC-stimulated transcytosis of fluorescent beads was lost and associated with a significant decrease in the number of glycoprotein-2 positive (Gp-2^+ve^) M cells. The results of these studies demonstrate that coronin 1a is required for uptake and transcytosis of luminal cargo across human M cells and that coronin 1a is necessary for differentiation of mature M cells that actively transcytose luminal gut antigens in response to pathogenic, but not commensal, microbes.

## INTRODUCTION

The most prominent feature of the follicle associated epithelium (FAE) is the presence of microfold (M) cells, which specialize in surveying the gut luminal environment by sampling and delivering luminal contents to immune cells as a part of normal intestinal homeostasis^1^. Luminal sampling by M cells is crucial for inducing protective mucosal immune responses but can also serve as the entry pathway for pathogens, including bacteria and viruses^2^. Enteric pathogens can influence the induction and/or function of intestinal M cells; however, the molecular mechanisms involved in the regulation of uptake and transcytosis of gut cargo by human M cells remains incomplete^3^. Although phagocytosis has been implicated as the entry pathway for luminal gut cargo into M cells, detailed mechanistic studies to determine the regulation of this pathway are also lacking. Following uptake, M cells move luminal antigens across the epithelium without being degraded, presumably since M cells have very few lysosomes^4^. The capacity of M cells to move cargo across the epithelium provides a route by which enteric pathogens can invade the intestinal mucosa. While numerous studies in animal models and cell lines have established that M cells are required for pathogen invasion of the Peyer’s patch, the mechanism responsible for trafficking gut cargo across human M cells has not been identified.

M cells are differentiated from actively dividing intestinal stem cells and their abundance in the FAE is affected by the presence and composition of intestinal microbiota as well as the presence of stromal and immune cells in the sub-epithelial dome of the Peyer’s patch. Compared to conventional mice, germ-free mice express very few M cells, yet exposure to a single enteric pathogen, such as Salmonella, can significantly increase M cell abundance, which occurs over many days^5–6^. Chronic gastrointestinal disorders such as ileal Crohn’s disease, which has a dysregulated microbiome as a component of disease, have increased numbers of M cells in the Peyer’s patch^7^. Additionally, short term exposure of the Peyer’s patch to pathogens increases the rate of M cell uptake of luminal antigens in the without altering M cell abundance^8^. These findings, among others, support the hypothesis that the M cell abundance and function are influenced by interactions with commensal or pathogenic organisms^9–10^. Considering the evidence that a relationship exists between the intestinal microbiome and mucosal immune responses, there is a need to identify the mechanisms responsible for how human M cells function in response to luminal challenges.

Understanding the factors that regulate human M cell expression and function requires a relevant human model. Previous studies have established protocols to differentiate human intestinal stem cell-derived enteroids (HIEs) to express functionally mature M cells. Recent reports by us, and others, demonstrate that M cell expressing HIEs can transcytose enteric pathogens, including Shigella and rotavirus^11,12^. However, our understanding of the mechanisms that regulate these processes in humans is incomplete. In the current study, we present a unique regulatory role for coronin 1a, a cytoskeletal remodeling protein not previously studied in the human intestinal epithelium, in the uptake and transcytosis of luminal cargo by human M cells.

## METHODS

### Antibodies/Reagents

Human recombinant TNF-α and RANK-L were from PeproTech. Mouse monoclonal antibody to glycoprotein-2 (Gp-2) was from MBL (Clone: 3G7-H9; Cat# D277-3). Mouse monoclonal antibodies to coronin 1a (Cat #: Ab56820) and β-actin (Cat #: 8224) were from Abcam. Rabbit polyclonal antibody to coronin 1a was from Abcam (HPA051132). Rabbit monoclonal antibody to CCL3 was from Abcam (Cat #: 170958). Fluorescent polystyrene latex beads (535/575 0.02-2.0μm), AlexaFluor 633-conjugated phalloidin, and Hoescht 33342 were from Life Technologies. Mouse monoclonal antibody to GAPDH was from Sigma (Cat #: G8795).

### Study approval

De-identified intestinal tissue was obtained from healthy subjects provided informed consent at Johns Hopkins University and all methods were carried out in accordance with approved guidelines and regulations. All experimental protocols were approved by the Johns Hopkins University Institutional Review Board (IRB# NA_00038329).

### Human Small Intestinal Enteroids (HIEs)

Human duodenal (n=1), jejunal (n=1), and ileal (n=4) enteroids were generated from biopsies obtained after endoscopic procedures utilizing the protocol established by the Clevers laboratory^13^. Enteroids were maintained as cysts embedded in Matrigel (Corning, USA) in non-differentiation media (NDM) containing Wnt3a, R-spondin-1, noggin and EGF, as we have described^14^. Enteroid monolayers (HEM) were generated as previously described in detail^14–15^. Monolayer differentiation was induced by incubation in Wnt3A-free and Rspo-1-free differentiation media (DFM) for five days^14^. M cell differentiation was accomplished in enteroid monolayers exposed basolaterally to DFM that included RANKL (100ng/ml) and TNF-α (50ng/ml) for 5 days (referred to as MCM)^16^. Monolayer confluency and differentiation were monitored by measuring transepithelial electrical resistance (TEER) with an ohmmeter (EVOM^2^; World Precision Instruments, USA), as previously described^14^.

### shRNA Lentivirus Transduction

Human ileal enteroids were transduced with GIPZ lentiviral shRNA kit for human Coronin 1a (GE Healthcare Dharmacon) based on protocol adapted from Heijmans et al.^17^. Coronin 1a shRNA 1 (V3LHS_412912: 5’-TAGTTTCTATATACAAGCA-3’), Coronin 1a shRNA 2 (V3LHS_412913: 5’-AACATGGGAAGTAACTCCT-3’), Coronin 1a shRNA 3 (V3LHS_412914: 5’-TCAACAAAAAGTACAACGT-3’). Following transduction, enteroids were maintained in NDM with 0.25μg/ml puromycin for stable selection of enteroids expressing each Coronin 1a shRNA clone. A separate enteroid line was also generated to stably express a non-silencing shRNA (NS) to serve as a negative control.

### Immunofluorescent Confocal Microscopy

Immunostaining was performed as previously described^14^. Briefly, ileal enteroid monolayers were grown to confluency on 24-Transwells (Corning 3470) and differentiated with MCM or DFM to express or lack Gp-2^+ve^ M cells, respectively. Monolayers were fixed in 4% paraformaldehyde (PFA) in PBS for 30 min prior to addition of blocking solution. Primary antibodies were exposed to permeabilized monolayers for 1 hour at RT and then overnight at 4°C. Cells were washed three times in PBS-T (PBS with 0.1% Tween-20) and exposed to secondary antibodies in the presence of phalloidin (AlexaFluor 633 conjugated; Life Technologies) and Hoescht 33342 (nuclear counterstain; Life Technologies) for 1 hour at RT. After washing in PBS-T, filters were removed from inserts and mounted (ProLong Gold Antifade Mountant, ThermoFisher) and coverslipped. Confocal images were obtained using Olympus FluoView 3000.

### M cell uptake and transcytosis

M cell mediated uptake was determined by confocal microscopy of apically exposed 2.0μm fluorescent polystyrene latex beads (Life Technologies; 4×10^8^ final concentration) to human enteroid monolayers expressing or lacking M cells that were cultured in the presence or absence of 10^6^ pathogenic bacteria overnight. Following infection, monolayers were washed, fixed, and stained for confocal microscopy. Uptake was defined as the number of internalized beads in Gp-2^+ve^ M cells and compared to the number of beads localized to the surface of the monolayers. Additionally, fluorescence intensity of internalized beads was quantified in cell lysates obtained following the overnight infection period. Cell lysate fluorescence was measured using Perkin Elmer Envision plate reader. M cell transcytosis of 0.02μm fluorescent beads was also quantified in basolateral media collected from treated monolayers using Perkin Elmer fluorescent plate reader.

### Western blot analysis

Cell lysates of human ileal enteroids were obtained by harvesting in lysis buffer (60 mM HEPES pH 7.4, 150 mM KCl, 5 mM Na_3_EDTA, 5 mM EGTA, 1 mM Na_3_VO_4_, 50 mM NaF, 1% Triton X-100) with protease inhibitor cocktail (1:100). Cells were disrupted by vigorous pipetting, flash frozen in liquid N2, and then end-over-end rotated for 2 hours at 4°C. Lysates were solubilized in sodium dodecyl sulfate gel-loading buffer with β-mercaptoethanol and heated at 70°C for 10 minutes. 30μg of total cell lystates of differentiated enteroids (with and without M cells) were separated on Tris-glycine gels and proteins transferred to nitrocellulose membranes. Protein expression was normalized to either GAPDH or β-actin as internal controls. Band intensity was detected and quantified using Li-Cor Odyssey Infrared Imaging System.

### Statistics

Data are shown as mean ± S.E.M. Statistical comparisons were performed in Prism 8 (GraphPad) using either the two-tailed Student’s t test or ANOVA with Tukey’s multiple-comparison post hoc test. P values less than 0.05 were considered significant. (\**p* < 0.05; \*\**p* < 0.01)

## RESULTS

### RANKL and TNF-α enhance the number of M cells in human ileal enteroid monolayers

Human enteroids can be differentiated to express M cells by reducing Wnt signaling and stimulating non-canonical NF-κB signaling by exposing enteroids to receptor activator of NF-κB ligand (RANKL)^11,16,18^. Stimulation of canonical and non-canonical NF-κB signaling enhances the expression of M cell associated genes in mice^19^ and the number of mature M cells in human intestinal epithelial models which mimics their abundance *in vivo*^11,16^. In humans, M cells comprise ~10% of the cells in the FAE^20^. We differentiated (no Wnt3a, no R-spondin-1; 100ng/ml RANKL, 50ng/ml TNF-α) human ileal enteroid monolayers (from established enteroid lines of 4 separate healthy donors) to express M cells^16^. Differentiating human ileal monolayers with both TNF-α and RANKL induced a significantly higher number of M cells compared to RANKL or TNF-α alone (**Figure 1A, 1B**). This finding is supported by a recent report showing similar effects in human enteroids following exposure to RANKL, retinoic acid, and lymphotoxin, suggesting that suppression of Wnt signaling and activation of NF-κB signaling enhances M cell expression in human intestinal enteroids (HIEs)^11^. We detected the M-cell specific marker, Gp-2, at the apical membrane of human enteroid monolayers (HEMs) (**Figure 1C**)^21,22^. Another hallmark of human M cells is the lack of apical microvilli^23,24^. To confirm that our M cell HEMs exhibit this morphology, we performed scanning electron microscopy and observed cells that lacked mature, densely packed microvilli and were surrounded by other epithelial cells with abundant microvilli (**Figure 1D**). We also show that the number of M cells in HEMs is higher than in duodenal and jejunal HEMs differentiated (DF) to express M cells (**Figure 1E**), further supporting the concept that HIEs recapitulate the phenotype of the intestinal segment from which they were derived^25^. Since our M cell differentiation media induced expression of Gp-2^+ve^ M cells with effaced apical membranes, we next tested whether our M cell model expresses human FAE markers^26^.

**Figure 1:**
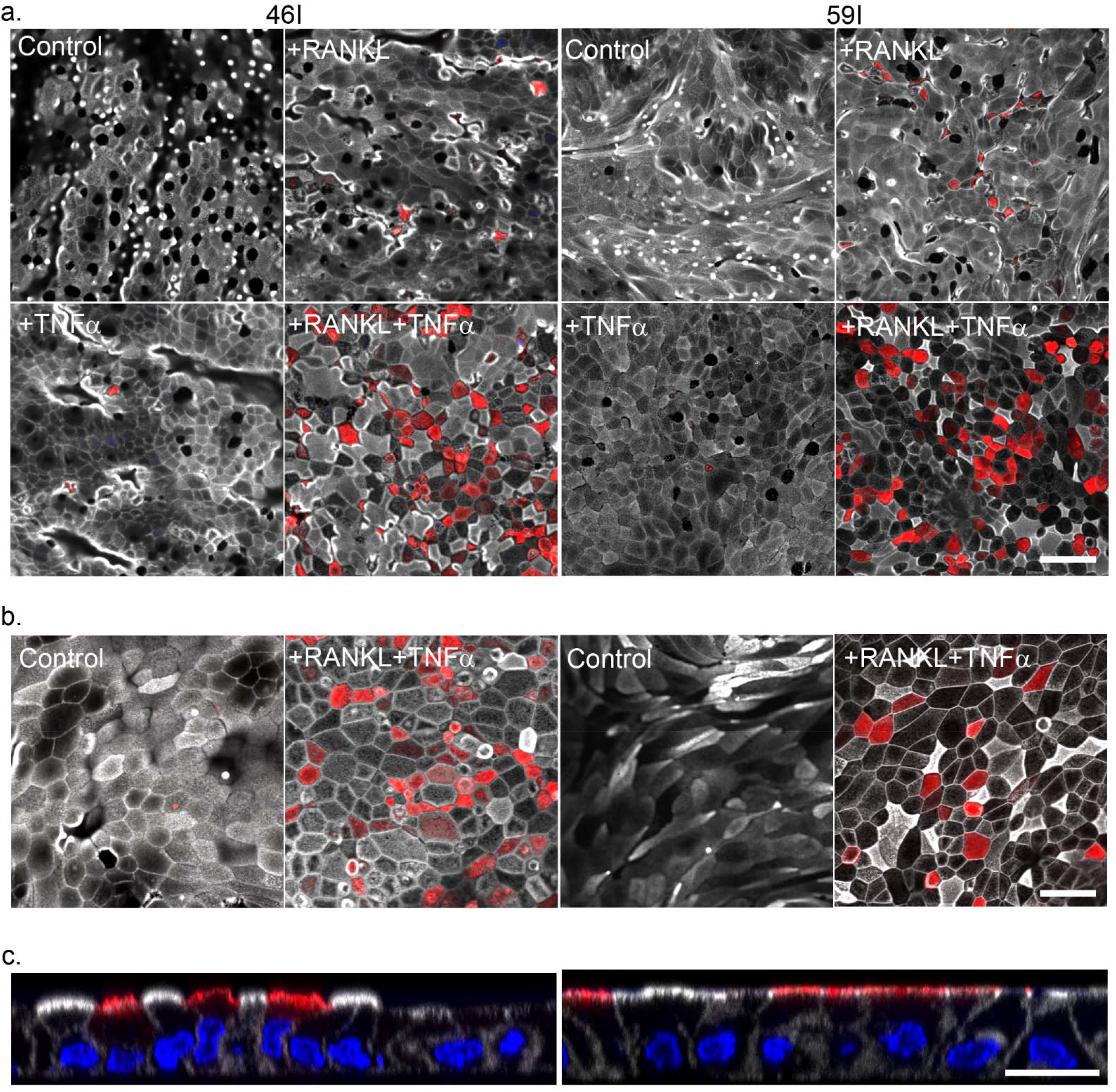
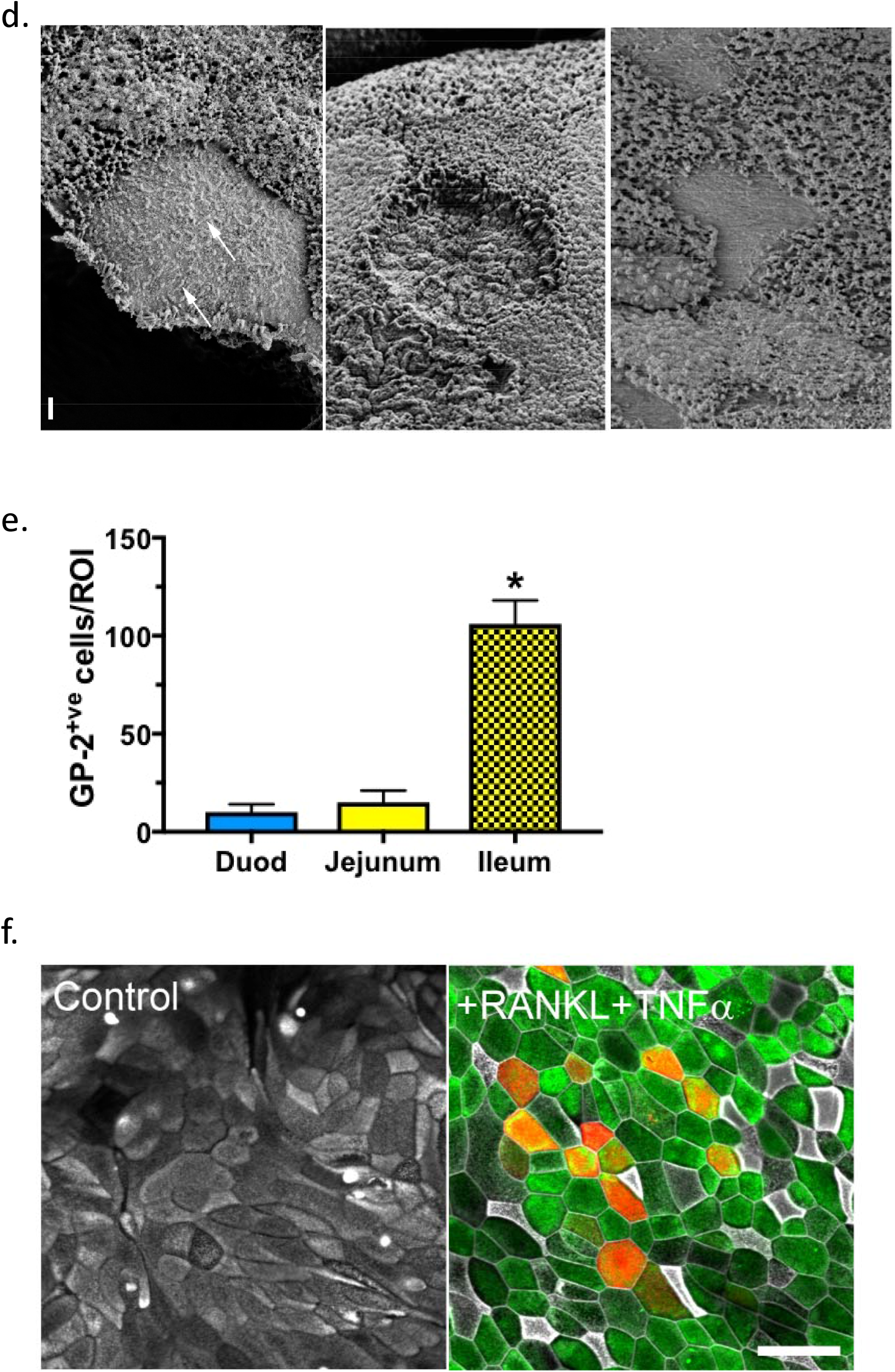
Induction of M cells in human enteroid monolayers. **A-C)** – Immunofluorescent confocal microscopy demonstrating the expression of Gp-2^+ve^ M cells (red) in ileal enteroid monolayers from four separate donors following differentiation in the presence or absence of RANKL and/or TNF-α. white=actin. **A** and **B** size bars = 50 μm. **C)** – Gp-2 expression was localized to the apical membrane of M cells HEMs. Red=Gp-2; white=actin; blue=nuclei. Size bar = 20 μm **D)** – Scanning electron microscopy of human ileal monolayers differentiated to express M cells. n=3. Size bar = 1.0 μm **E)** – M cells abundance is highest in ileal enteroid monolayers differentiated with RANKL and TNF-α compared to duodenal (Duod) and jejunal enteroid monolayers. M cell numbers were quantitated from at least 10 ROIs (regions of interest). Results are mean ± SE from 3 independent experiments. *p<0.05. **F)** – Confocal microscopy shows CCL23 (green) in cells that lack Gp-2 (red) suggesting that M cell enteroids also express FAE markers. Blue = nuclei; white = actin. Size bars = 50μm

### Differentiation with RANKL and TNF-a induces expression of the FAE marker, CCL23

The FAE is the epithelial “dome” of the Peyer’s patch that covers the underlying lymphoid follicle and expresses specific chemokines and pattern recognition receptors while lacking secretory cells and expression of digestive enzymes^27,28^. This pattern of protein expression and lineage differentiation are distinct from the small intestinal villus epithelium. To better characterize the profile of our M cell HEMs, we determined whether M cell HEMs express known markers of the FAE and M cells^2^ including transcription factors SOX8, Spi-B, and chemokines. For example, we validated the expression of CCL23, a chemokine expressed in the human FAE^26^, in M cell HEMs compared to DF enteroids, which lacked CCL23 expression (**Figure 1F**). Similar findings were recently observed in human enteroids expressing M cells by RNAseq^11^. In this study, we demonstrate in M cell expressing HEMs, nearly every cell, with the exception of Gp-2^+ve^ M cells, expressed CCL23. These data demonstrate that differentiating ileal HEMs to express M cells also induces expression of non-M cell FAE markers and thus is a relevant *ex vivo* model of the human FAE.

### Enteric pathogens induce M cell-mediated uptake and transcytosis

M cells sample luminal antigens across the follicle associated epithelium for delivery to underlying innate and adaptive immune cells; however, how this process is regulated in human M cells remains to be elucidated. We measured human M cell function (i.e. uptake and transcytosis) and compared them to DF (i.e. “villus”-like) HEMs. DF or M cell HEMs, were exposed to fluorescent polystyrene latex beads (**Figure 2A-E**), a standard assay performed in numerous published reports to measure M cell uptake and transcytosis^11,18,29–31^. Under basal conditions, apical entry and transcytosis of fluorescent beads (detected in the basolateral compartment of HEMs) was similar between DF and M cell monolayers (black bars; **Figure 2B**). We challenged DF and M cell HEMs to *Enteropathogenic E. coli* (EPEC), which binds the surface of human M cells^32^. Exposure of M cell enteroids to live EPEC (blue bars) significantly increased bead uptake and transcytosis (**Figure 2B**). Similar results were observed in M cell HEMs exposed to sonicated EPEC (yellow bars) but not lipopolysaccharide (LPS; white bars). EPEC did not induce transcytosis in DF HEMs. These findings suggest that outer membrane proteins or secreted factors from EPEC are required for luminal bead capture and transcytosis by M cells. Since our EPEC data suggest that pathogenic microbes can increase bead capture, we performed similar studies with the commensal bacterial strain, HS. Apical exposure of HS did not increase bead uptake or transcytosis compared to vehicle control and EPEC infected M cell HEMs (**Figure 2C**). These results suggest that commensal bacteria do not stimulate M cell function, which instead requires factors from enteric pathogens. A similar effect was observed in M cell HEMs exposed to the invasive AIEC strain (**Figure 2D**). In this experiment, AIEC induced internalization of fluorescent beads into Gp-2^+ve^ M cells while the beads remained in the luminal compartment in normally differentiated (DF) HEMs or in unexposed M cell HEMs (**Figure 2E**).

**Figure 2:**
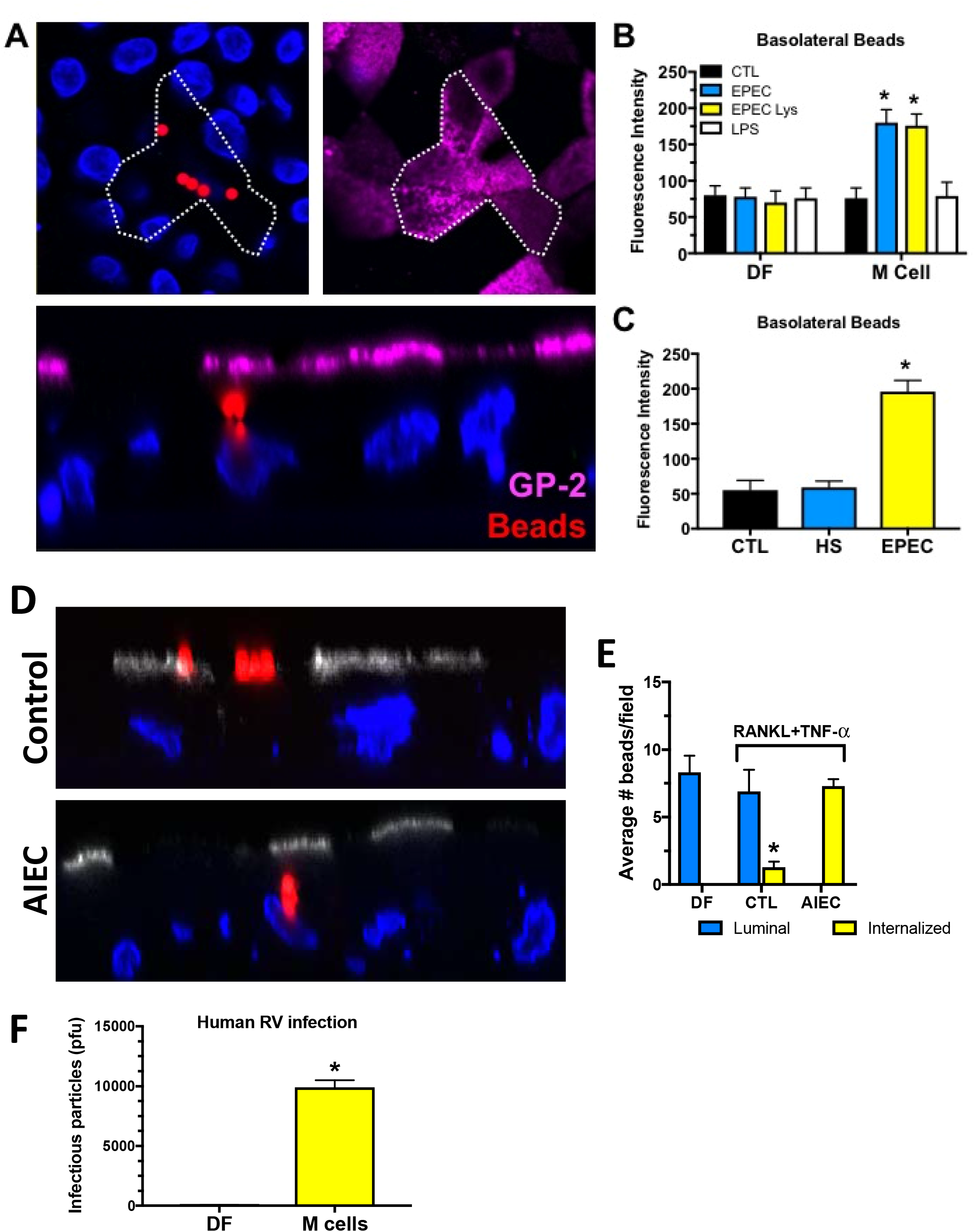
Luminal capture of fluorescent beads in human ileal M cell expressing monolayers is stimulated by EPEC or AIEC, not commensal bacteria, a process that does not involve LPS. **A)** Immunoflourescence confocal microscopy of M cell expressing enteroid monolayers exposed to 2.0μm fluorescent polystyrene latex beads (red). Beads were only detected in Gp-2 (magenta) expressing M cells. **B)** Fluorescent beads (2.0μm) were added apically to human enteroid monolayers and exposed to vehicle (CTL), LPS, intact EPEC, or sonicated EPEC (EPEC lys). Monolayers were lysed and the number of internalized beads was measured in lysates using a fluorescent plate reader. **C)** Fluorescent beads (0.02μm) were added apically to human monolayers and exposed to vehicle (CTL), the commensal bacteria, HS, or EPEC. Basolateral media was collected and fluorescence intensity of beads measured on a plate reader. Results are mean ± SE from 3 independent experiments. *p<0.05. **D)** Immunofluorescence confocal microscopy of M cell HEMs either mock infected (Control) or exposed to AIEC (Strain 857c, 10^6^ CFUs for 16h). Approximately 4×10^8^ fluorescent polystyrene latex beads (red) was added apically with or without AIEC to visualize uptake by M cells, labeled with anti-Gp-2 antibody (white). Blue = nuclei. **E)** AIEC-induced uptake of fluorescent beads was quantified in human M cells. In DF and uninfected (CTL) M cell HEMs, fluorescent beads were observed only on the apical (luminal) surface. Following AIEC exposure, M cell HEMs internalized all fluorescent beads. Results are mean ± SE from 3 independent experiments. *p<0.05. **F)** Transcytosis of human rotavirus (RV) requires M cells. Human RV strain, Ito, was exposed apically (24h) to DF or M cell enteroid monolayers. Infectious virus was quantified from basolateral media by plaque assay. pfu = plaque forming units. Results are mean ± SE from 3 experiments. *p<0.01

We also tested whether enteric viruses transcytose across M cell enteroids by evaluating human rotavirus (RV). Transcytosis of human RV (Ito strain^33^) occurred in M cell HEMs but not in DF HEMs (**Figure 2F**). These results mimic similar findings recently reported in human M cell monolayers^11^. Our data support the concept that luminal capture and transcytosis can be initiated by pathogenic microbes, including viruses, in human M cell enteroids.

### Coronin 1a is necessary for uptake of luminal cargo by human M cell expressing enteroids

We next asked what protein mediates the M cell response to enteric pathogens. Since human M cell HEMs demonstrate increased uptake and transcytosis of luminal cargo when exposed to enteric pathogens, we hypothesized that coronin 1a may regulate this process. A role for coronin 1a in pathogen induced activation of phagocytosis has been described^34–36^. Coronin 1a mediates phagocytic entry and protection of pathogenic, but not commensal, mycobacteria from lysosomal degradation in macrophages^34^; however, there have been no studies to understand the role of coronin 1a in the human intestinal epithelium. Since sampling of the luminal gut environment occurs via phagocytosis in M cells, we asked whether differentiation of human enteroid monolayers to express M cells also stimulated expression of phagocytic proteins such as coronin 1a. As shown in **Figure 3A**, coronin 1a protein expression is increased in differentiated HEMs following RANKL treatment; however, this induction was greater when exposed to both RANKL and TNF-α. Coronin 1a expression was increased >20- fold in M cell HIEs compared to DF HIEs, which we validated by immunoblot (**Figure 3A**). Confocal microscopy showed that Gp-2^+ve^ M cells co-localized with coronin 1a (**Figure 3B**). Analysis of confocal images confirmed that coronin 1a expression was highly correlated with M cells (r^2^=0.9922; **Figure 3C**) while only ~30% of coronin 1a expressing cells were also positive for Gp-2 (r^2^=0.0981; **Figure 3D**). In our M cell expressing HEMs, the number of coronin 1a expressing cells was nearly 3 times higher than Gp-2^+ve^ mature M cells (**Figure 3E**). The distribution of coronin 1a included expression at the apical pole of epithelial cells which was also associated with actin filaments along the lateral membrane (**Figure 3F**) supporting its already known role as an cytoskeletal protein.

**Figure 3:**
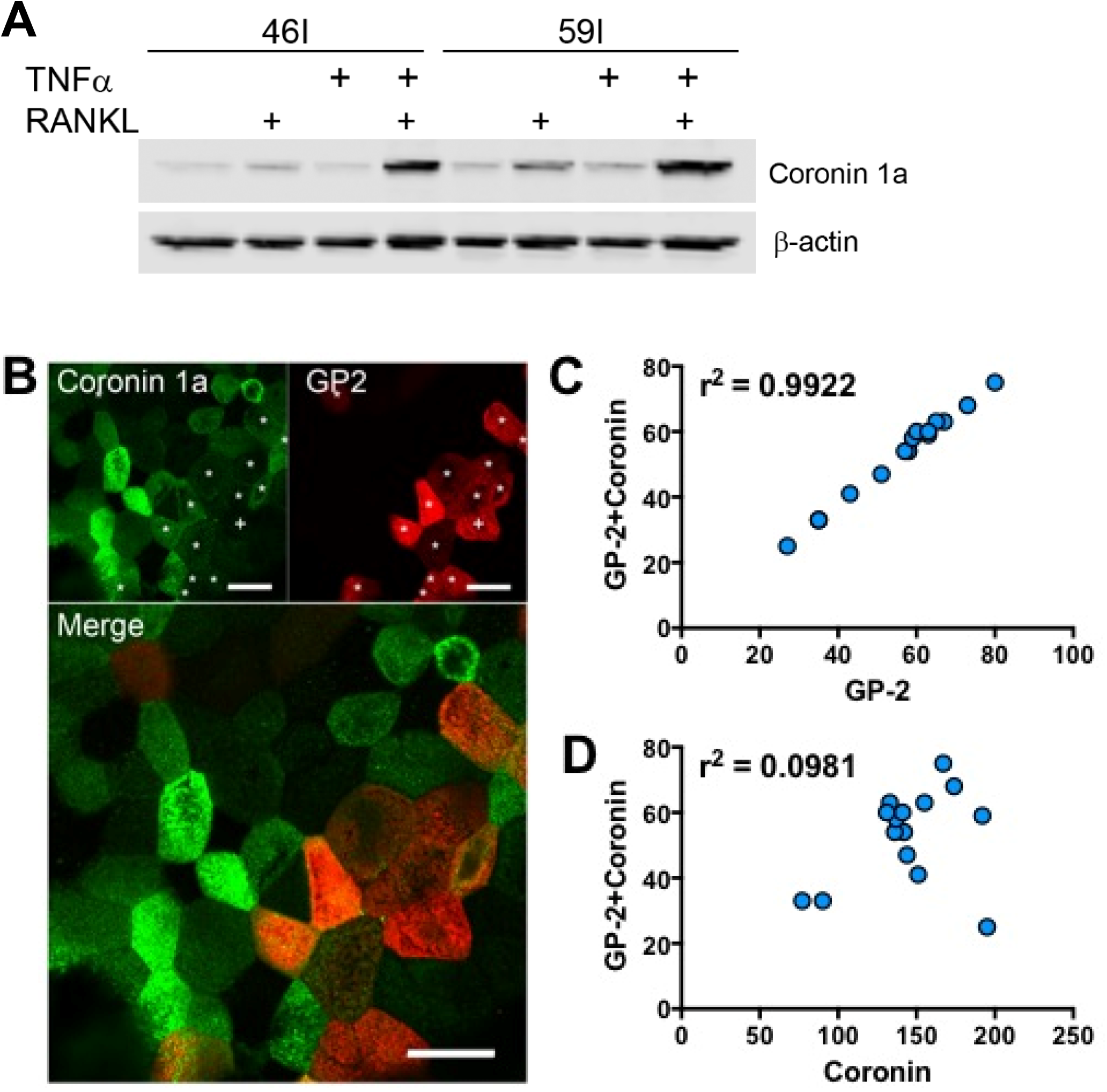
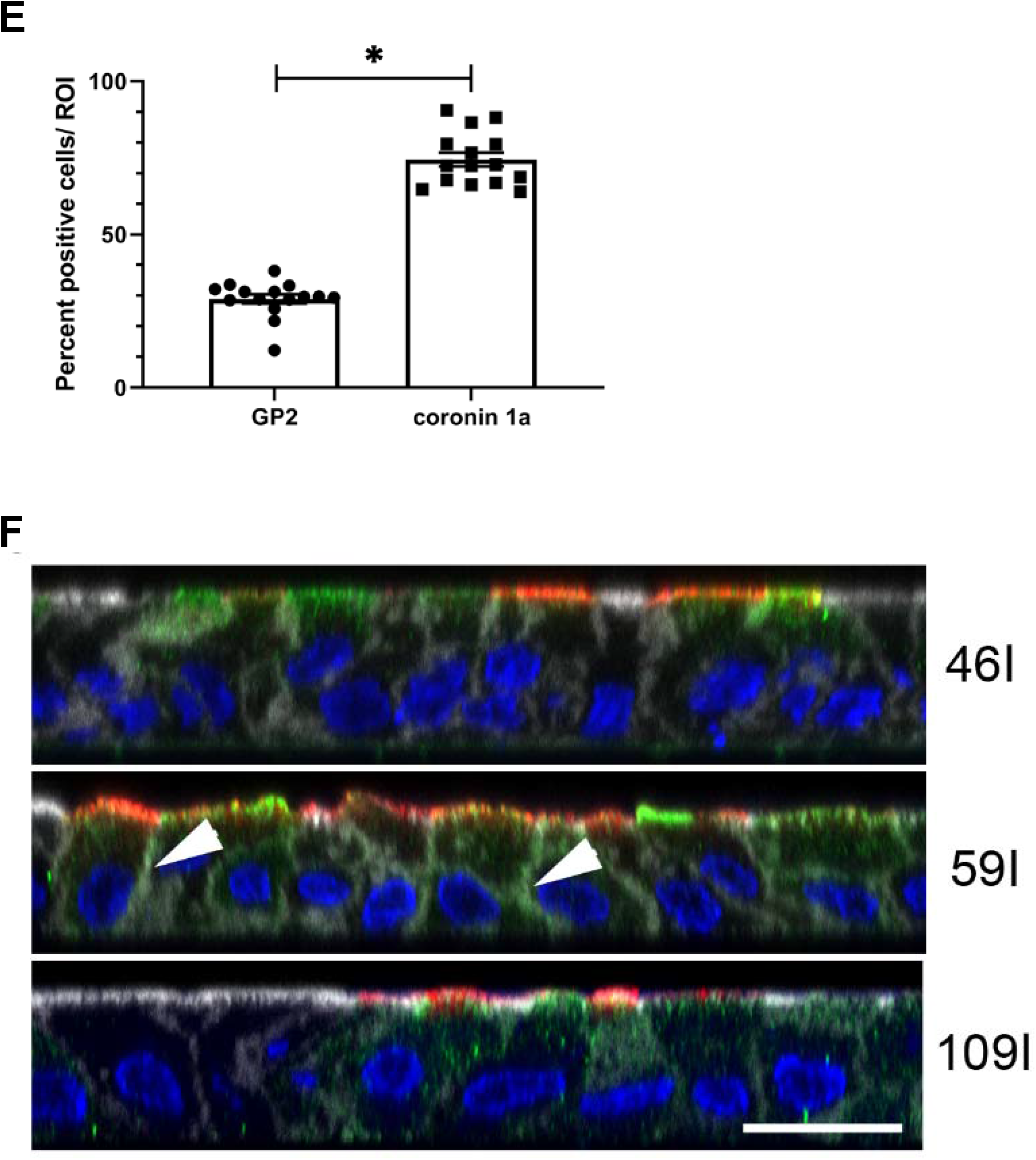
Coronin 1a expression highly correlates with Gp-2 expressing M cells. **A)** M cell differentiation with RANKL + TNF-α induced the highest expression of coronin 1a. **B)** Coronin 1a (green) and Gp-2 (red) co-localize in M cell enteroid monolayers. Asterisks (white) indicate co-expressing cells. Size bars = 20 μm **C)** Pearson’s correlation analysis for co-localization of coronin 1a and Gp-2 demonstrated a near perfect correlation between M cells and coronin 1a while **(D)** only ~30% of coronin 1a expressing cells were also positive for Gp-2. Data are correlation coefficients from n=3 separate enteroid lines. **E**) M cell differentiation with RANKL + TNF-α induced the highest expression of coronin 1a. **F**) Confocal microscopy of HEMs expressing coronin 1a (green) and Gp-2 (red). XZ projections show apical Gp-2 with coronin 1a at the apical domain of the epithelium and partial co-localization (arrowheads) with actin (white) at the lateral membrane. N=3 separate enteroid lines. Size bar = 20 μm.

To demonstrate a role for coronin 1a in human M cell function, we generated a stable coronin 1a knockdown (KD) ileal enteroid line (shRNA construct 59; ~70% decreased protein expression) as well as a control HIE in which coronin 1a expression was not affected by shRNA KD constructs (NS) (**Figure 4A**). As demonstrated in **Figure 4B**, shRNA construct 59 KD of coronin 1a decreased protein expression in HEMs differentiated with RANKL and TNF-α to express mature M cells when compared to the NS control enteroid line. Using these lines, we investigated the role of coronin 1a in mediating pathogen-induced activation of M cell uptake and transcytosis.

**Figure 4:**
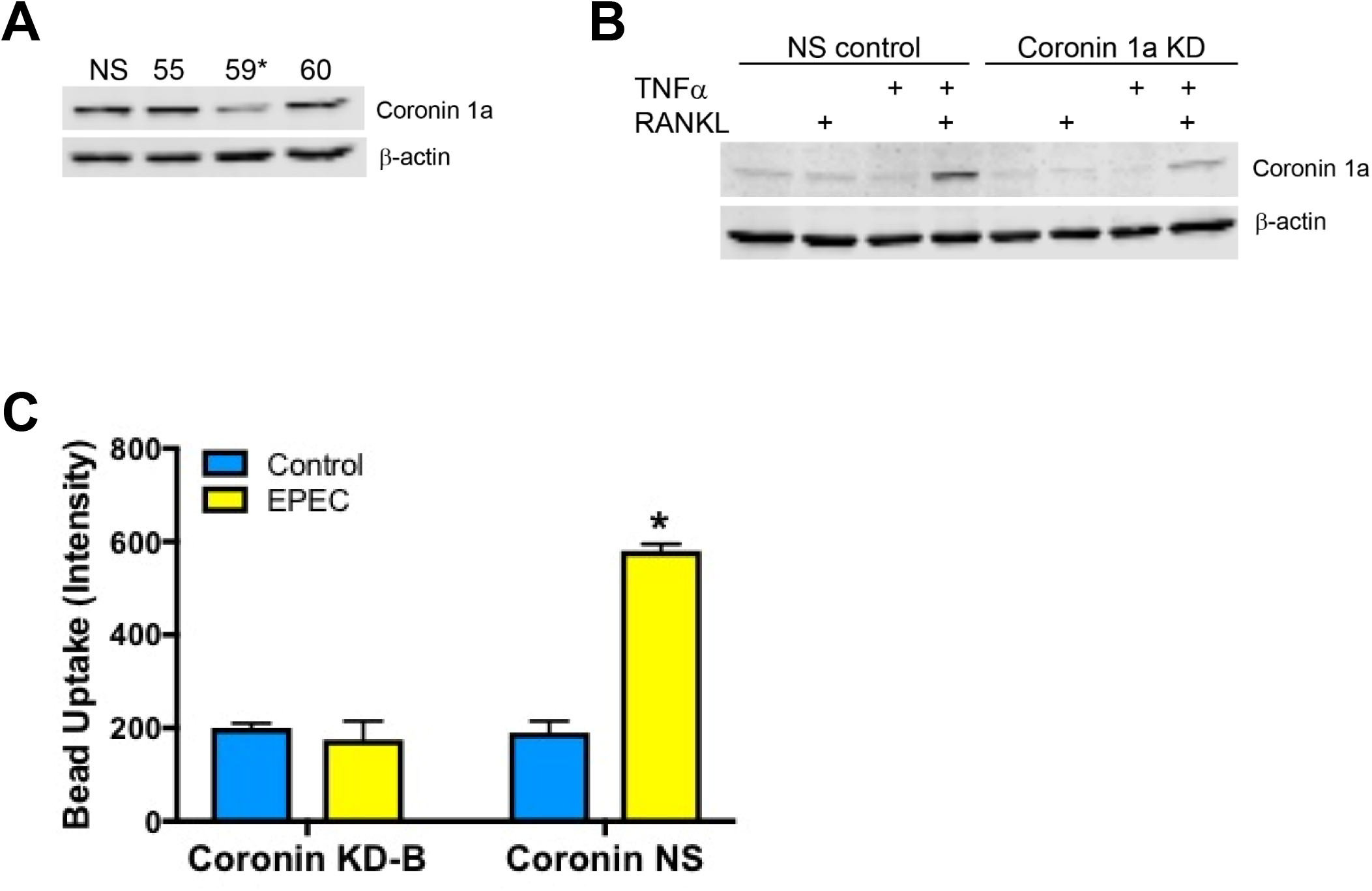
Luminal capture and transcytosis by M cell expressing human ileal enteroid monolayers requires coronin 1a. **A-B)** Western blots of HIEs transduced with coronin 1a shRNA lentiviruses. The #59 construct has most reduced coronin 1a expression compared to non-silencing (NS) and non-transduced wild type controls (data not shown). **C**) Coronin 1 knockdown (KD-B) prevented EPEC mediated luminal uptake of fluorescent beads. Basolateral media was collected and fluorescence measured on a plate reader. Results are mean ± SE from 3 independent experiments. *p<0.01.

To test the possibility that that pathogens may exploit coronin 1a to facilitate capture and passage of gut contents across the intestinal epithelium, we performed luminal bead capture studies in coronin 1a KD enteroids exposed to EPEC. Coronin 1a KD enteroids lost their ability to uptake fluorescent beads compared to control enteroid monolayers (NS) (**Figure 4C**) suggesting that coronin 1a is required for entry of non-specific cargo into human M cells. These data demonstrate that EPEC can induce capture and transcytosis of luminal content across human M cell enteroid monolayers by a coronin 1a-dependent mechanism.

### Coronin 1a is required for expression of Gp-2^+ve^ M cells

M cells are differentiated from Lgr5^+ve^ intestinal stem cells (ISCs)^29^ via RANK signaling, which is responsible for lineage differentiation of M cells^37^. RANKL induces the expression of M cell-specific transcription factors including SOX8 and Spi-B, which are essential for differentiation of M cells^29,37–40^. Mature M cells are defined by the expression of the pattern recognition receptor (PRR), glycoprotein 2 (Gp-2), at the apical membrane^21,22^. In addition to Gp-2, which binds FimH of *E. coli,* M cells specifically express other PRRs that bind and transport microbes across the epithelium^41^; however, what regulates apical PRR localization in M cells is not known. Since our results suggest that the cytoskeletal remodeling protein, coronin 1a, is required for pathogen-induced stimulation of M cell function and we found a high correlation between Gp-2^+ve^ cells and coronin 1a, we hypothesized that loss of coronin 1a affects Gp-2 expression and/or localization and could explain why EPEC is unable to induce M cell activity in the absence of coronin 1a. As shown by confocal microscopy in **Figure 5A**, KD of coronin 1a resulted in decreased abundance of Gp-2^+ve^ M cells in HEMs. Quantification of the number of Gp-2^+ve^ M cells in coronin 1a-KD enteroid lines was decreased by >90% when compared to non-silenced (NS) control HEMs or non-transduced HEMs that were differentiated to express M cells (**Figure 5B**). These data suggest a critical role for coronin 1a in regulating the maturation of Gp-2^+ve^ human M cells.

**Figure 5:**
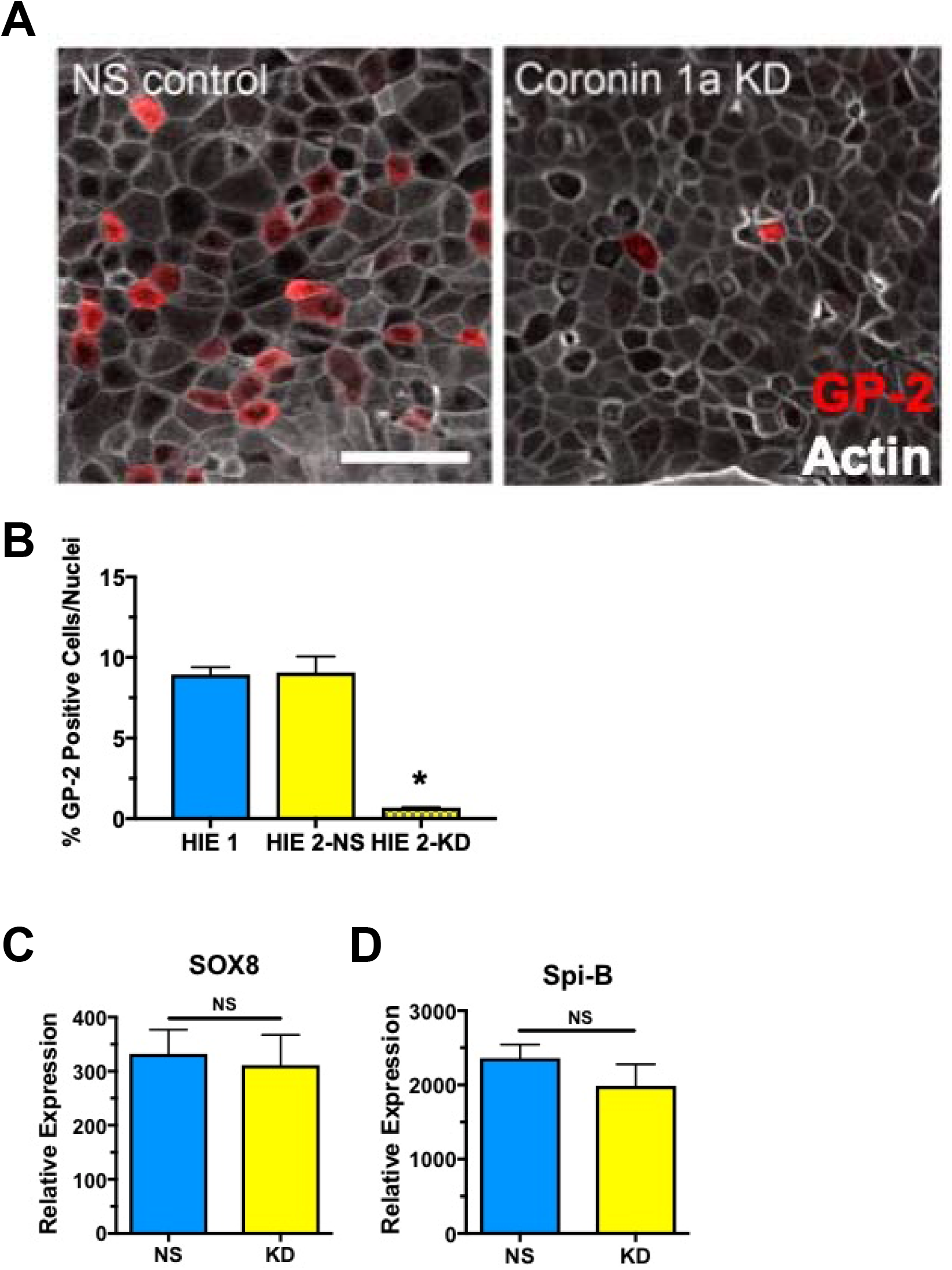
Coronin 1a is necessary for the expression of mature M cells in human ileal enteroid monolayers. **A)** Human ileal M cell monolayers expressing (NS control) or lacking (KD) coronin 1a were differentiated to express M cells and stained for expression of Gp-2 (red). Size bar = 50 μm **B)** Coronin 1a KD (sh-B) resulted in significant decrease in the number of Gp- 2^+ve^ M cells when compared to non-silencing controls (NS) from the same patient line (HIE-2) and a separate normal, ileal enteroid monolayer (HIE-1). Results are mean ± SE from 3 independent experiments. *p<0.01. **C-D**) Ileal HEMs were differentiated with RANKL+TNF-α to express M cells in coronin NS (non-silencing) and KD lines. Decreased coronin 1a expression did not affect mRNA expression of transcription factors, SOX8 or Spi-B. Data are mean ± SE from n=3 independent experiments. All results were not statistically different.

In order to demonstrate that loss of coronin 1a results in decreased apical expression of the PRR, Gp-2, and not a loss of M cells, we determined the expression of the transcription factors SOX8 and Spi-B in coronin 1a KD HEMs. We performed quantitative PCR (qPCR) in NS and coronin 1a KD HEMs and observed no difference in the mRNA levels of either SOX8 (**Figure 5C**) or Spi-B (**Figure 5D**). Since SOX8 and Spi-B are necessary for differentiation of M cells, our data suggest that the mature M cell phenotype (i.e. Gp-2^+ve^ expression) is dependent on coronin 1a expression.

## DISCUSSION

The results of the current study are the first to characterize a role for coronin 1a in regulating the function of human M cells in a primary human intestinal epithelial model (i.e. enteroids). Moreover, we demonstrate in this model that human M cells functionally respond to luminal signals that induce transcytosis of luminal gut cargo. We show that enteric pathogens, including EPEC, AIEC, and rotavirus, but not commensal bacteria, are sufficient to stimulate M cell-mediated uptake and transcytosis of luminal contents. In the case of EPEC, we show that either intact or sonicated EPEC can initiate an M cell response, but this effect does not involve TLR4 as LPS does not mimic the same effect. In order to understand what M cell related protein is responsible for this pathogen-induced stimulation of M cell function, we revealed a unique regulatory role for coronin 1a. In HEMs enriched for M cells, coronin 1a demonstrates a high correlation of expression with Gp-2^+ve^ mature M cells. Using coronin 1a KD HEMs, we discovered that EPEC induction of M cell activity was lost. This finding was associated with decreased abundance of Gp-2^+ve^ in M cells; although, expression of the M cell transcription factors, SOX8 and Spi-B, were unaffected.

To better understand the mechanisms that govern human M cell function, we employed the use of human enteroids derived the ileum of healthy donors. Human enteroids are differentiated to express M cells by reducing Wnt signaling and stimulating non-canonical NF-κB signaling by exposing enteroids to receptor activator of <u>N</u>F-<u>κ</u>B <u>l</u>igand (RANKL)^11,16,18,19,29^. Recent reports have shown that in addition to RANKL, stimulation of canonical TNFR-mediated NF-κB signaling results in increased numbers of M cells in human enteroids that better mimics their abundance in vivo^11,16,20^. This additive activation of NF-κB signaling has been shown to significantly enhance the expression of M cell associated genes in mice^19^ but without affecting the number of Gp-2^+ve^ M cells *in vivo* and in murine enteroids. For the current study, we differentiated (no Wnt3a, no R-spondin-1; 100ng/ml RANKL, 50ng/ml TNF-α) human ileal enteroid monolayers to express M cells and demonstrate that both TNF-α and RANKL induce a significantly higher number of M cells compared to RANKL alone. This finding is supported by a recent report showing similar effects in human enteroids following exposure to RANKL, retinoic acid, and lymphotoxin suggesting that concomitant suppression of Wnt signaling and NF-κB activation converging at RelB enhances M cell expression in HIEs^11^. To further support the relevance of human M cell expressing HEMs, we show that M cell abundance is highest in ileal enteroids compared to those from the duodenum or jejunum. Since Peyer’s patches are more prevalent in the distal small intestine, our data supports previous findings that HIEs can recapitulate the phenotype of the intestinal segment from which they were derived^25^.

Coronin 1a plays a role in phagocytosis of pathogenic bacteria by antigen presenting immune cells^34^. To date, a role for coronin 1a in human epithelial cells has not been described since studies designed to interrogate an epithelial-specific role for coronin 1a have not been performed. Our data are the first to identify that coronin 1a is expressed in primary human intestinal epithelial cells modeled to mimic the follicle associated epithelium (FAE) of the Peyer’s patch. While RANKL induced some increased expression of coronin 1a in HEMs, this expression was greatly enhanced when both TNF-α and RANKL were added to induce M cell differentiation. This enhanced coronin 1a expression was also reported in a recent report from Ding et al.^11^ which found similar upregulation of coronin 1a by RNAseq in human ileal enteroids differentiated with RANKL, retinoic acid, and lymphotoxin. These data, together with ours, suggest that stimulation of canonical and non-canonical NF-κB signaling is sufficient to increase coronin 1a in human enteroids expressing M cells. Considering that the FAE only comprises a fraction of the entire intestinal epithelium could explain how epithelial expression of coronin 1a could be overlooked. Interestingly, analysis of large data sets of murine Peyer’s patches or murine enteroids differentiated to express M cells did not reveal enhanced expression of coronin 1a. Whether coronin 1a plays a similar role in regulating M cell function in the murine FAE remains to be determined and would be important to reveal whether regulation of M cell activity in humans and mice differ. Additionally, since M cells are present in other mucosal surfaces, including the lung and nasal epithelium, further studies investigating a role for coronin 1a in these tissues would also be of interest.

While our findings strongly suggest that lineage differentiation of human M cells is sufficient to express coronin 1a, which is necessary to mediate luminal responses and facilitate M cell function, other factors with the Peyer’s patch are known to influence M cell activity. Evidence in mouse studies show that stromal and immune cell derived factors, such as 4-1BB and S100A4, from the sub-epithelial dome (i.e. stromal cells immediately below the FAE) can enhance M cell function without affecting the number of M cells^30,42^. Whether these factors have the same effects on human M cells remains to be determined. In addition to receiving signals, intestinal epithelial cells secrete cytokines/chemokines that instruct immune responses to infections. Similarly, M cells associate with a sub-type of intestinal macrophages and dendritic cells that are phenotypically distinct from the same monocyte-derived immune cells present in intestinal villi immediately adjacent to the PP^43^. These findings suggest that factors from other stromal cells, which are not present in our current M cell expressing HEMs, may influence M cell function. Whether coronin 1a maintains a similar regulatory role in the presence of other stromal regulatory factors is yet unknown. Since co-cultures of human enteroids with stromal, endothelial, and immune cells have already been established^14,44,45^, future studies using these more complex in vitro cultures could provide additional insight in human M cell function.

Our results demonstrate the requirement for coronin 1a in regulating M cell function during infection with pathogenic microbes; however, the signaling mechanisms initiated by pathogenic bacteria that lead to activation of coronin 1a remain to be studied in human M cells. Coronin 1a activity is regulated by Ser/Thr phosphorylation by PKC^46^, which is also associated with elevation of intracellular calcium^47^. Whether AIEC, EPEC, and RV, similarly or differentially affect the expression and/or activation of coronin 1a in human M cells requires deeper investigation and is beyond the scope of the current study. In addition to phosphorylation, our data demonstrate that coronin 1a is critical for M cell-mediated antigen uptake and transcytosis in response to an enteric pathogen. It is unclear what mechanism explains this loss of uptake and transcytosis in human M cells when coronin 1a expression is reduced. A role for coronin 1a in the anchoring, retention, or trafficking of apical proteins remains to be investigated, particularly in human M cells. Considering our data showing that Gp- 2 expression is significantly diminished in coronin 1a KD HEMs, it is reasonable to hypothesize that coronin 1a may stabilize the apical expression of PRRs in M cells. Future studies to investigate whether coronin 1a affects other PRRs such as β1-integrin (binds *Yersinia pseudotuberculosis*^48^) and dectin-1 (binds secretory IgA^49^ and fungal β-glucan^50^) in our human M cell HEMs would provide more insight into the regulation of human M cell function by coronin 1a.

Our studies are the first to demonstrate that human M cell expressing enteroids respond to signals from luminal pathogenic microbes to induce phagocytic uptake and transcytosis of gut cargo by a unique coronin 1a dependent mechanism. The results of these current study will improve our understanding how human M cells function for future development of novel therapeutic strategies to treat GI disorders, such as Crohn’s disease, or even possibly development of new strategies to enhance oral vaccine delivery.

## Acknowledgments

The authors wish to thank Dr. Alfredo Torres (University of Texas Medical Branch) for providing the AIEC strain NRG857c (O83:H1). The authors also acknowledge the Integrated Physiology and Imaging Cores of the Hopkins Conte Digestive Disease Basic and Translational Research Core Center. This project was funded by NIH/NIAID P01-AI125181 and NIH/NIDDK P30- DK089502.

